# Micro-Magellan: A flexible, open source acquisition software for high throughput biological light microscopy

**DOI:** 10.1101/038117

**Authors:** Henry Pinkard, Nico Stuurman, Kaitlin Corbin, Ronald Vale, Matthew F. Krummel

**Author notes:** Corresponding Author: Henry Pinkard.

## Abstract

We demonstrate the capabilities of μMagellan: a flexible, open source microscopy software for reproducible high throughput imaging of biological samples across heterogeneous scales of space and time. μMagellan provides a simple user interface for exploration and automated imaging of non-cuboidal regions. By utilizing the hardware abstraction layer of μMagellan, μMagellan provides a powerful and extensible platform for imaging heterogeneous biological samples on a wide range of existing microscopes.

---

The past decade has seen an explosion of innovative techniques in optical microscopy. The potential of these techniques to reveal the complex orchestra of biological processes across large scales of space and time has increasingly been recognized.^1,2^ Two barriers to fully realizing the potential of any new optical techniques are instrumentation capable of acquiring the requisite data and software capable of analyzing it. The latter need has been met by the development of several software packages for the visualization and analysis of terabyte sized imaging volumes.^3-5^ Widefield, confocal, and multiphoton microscopes already present in many labs provide the requisite hardware for high throughput microscopy. However, the full potential of these instruments is limited in many cases by the lack of automated and customizable software.^6^

Biological specimens come in unusual shapes, and the spatial organization of the cells and structures within the specimen is frequently unknown prior to a period of manual exploration. Thus, there is a critical need for software that allows for efficient and intuitive navigation of samples. In addition, the vast majority of biological processes do not occur in cuboidal imaging volumes, and acquiring empty or irrelevant space limits the scalability of high-throughput imaging approaches. A more flexible, specimen-adaptive acquisition concept is needed to enable systematic, quantitative imaging of biological samples.^1^ Finally, reproducibility is an essential component of such approaches. Reproducibility in biological imaging requires the association of instrument settings with the common morphological features of specimens across samples and therefor the integration of that morphological information into acquisitions. Software packages have begun to address a subset of these challenges, but not without requirements for specific hardware or in house programming expertise to adapt them to different hardware platforms.^4^

Here we present μMagellan, a flexible and powerful software platform for enabling high throughput 2D and 3D data collection across a range of existing microscopy modalities. μMagellan utilizes the hardware abstraction layer of μMagellan^7,8^ allowing it to be used with a diverse set of components or complete microscopes from different vendors. μMagellan provides several key features for imaging biological systems at a variety of scales of space and time, the combination of which dramatically reduces the time and effort researchers must expend to perform experiments. μMagellan is designed according to the best practices of bioimaging software usability^9^, including in-application documentation, an intuitive user interface, cross-platform usability, open source code, and scalability across space and time. In addition, image data written by μMagellan can be opened directly in the BigDataViewer^3^, a FIJI^10^ plug-in designed for the visualization and analysis of large volumetric imaging data. BigDataViewer provides virtualized pixel access to an intelligent rendering scheme that can re-slice data from arbitrary angles for interactive visualization, and integration with FIJI and the image processing library ImgLib2^11^ provides access to a library of existing image processing algorithms including those for filtering segmentation, and deconvolution.

A common challenge when dealing with large 3D samples in microscopy is visualizing and navigating structure at a variety of scales to find areas of interest.^12^ To address this, μMagellan provides “explore acquisitions”, which allow users to navigate samples in 3D space, and collect tiled, high-resolution areas of planar images or z-stacks with a single click **(Fig. 1, Supplementary Note 1).** This enables rapid mapping of areas of interest within samples while giving users complete control to minimize photobleaching. Exploration creates and stores a multi-scale map of samples, from which subsequent imaging can be precisely specified. The acquisition UI enables panning and zooming through 2D slices of samples or arbitrary size, thanks to a multiresolution file format **(Supplementary note 2)** that allows random access to pixel data at any resolution.

**Figure 1.**
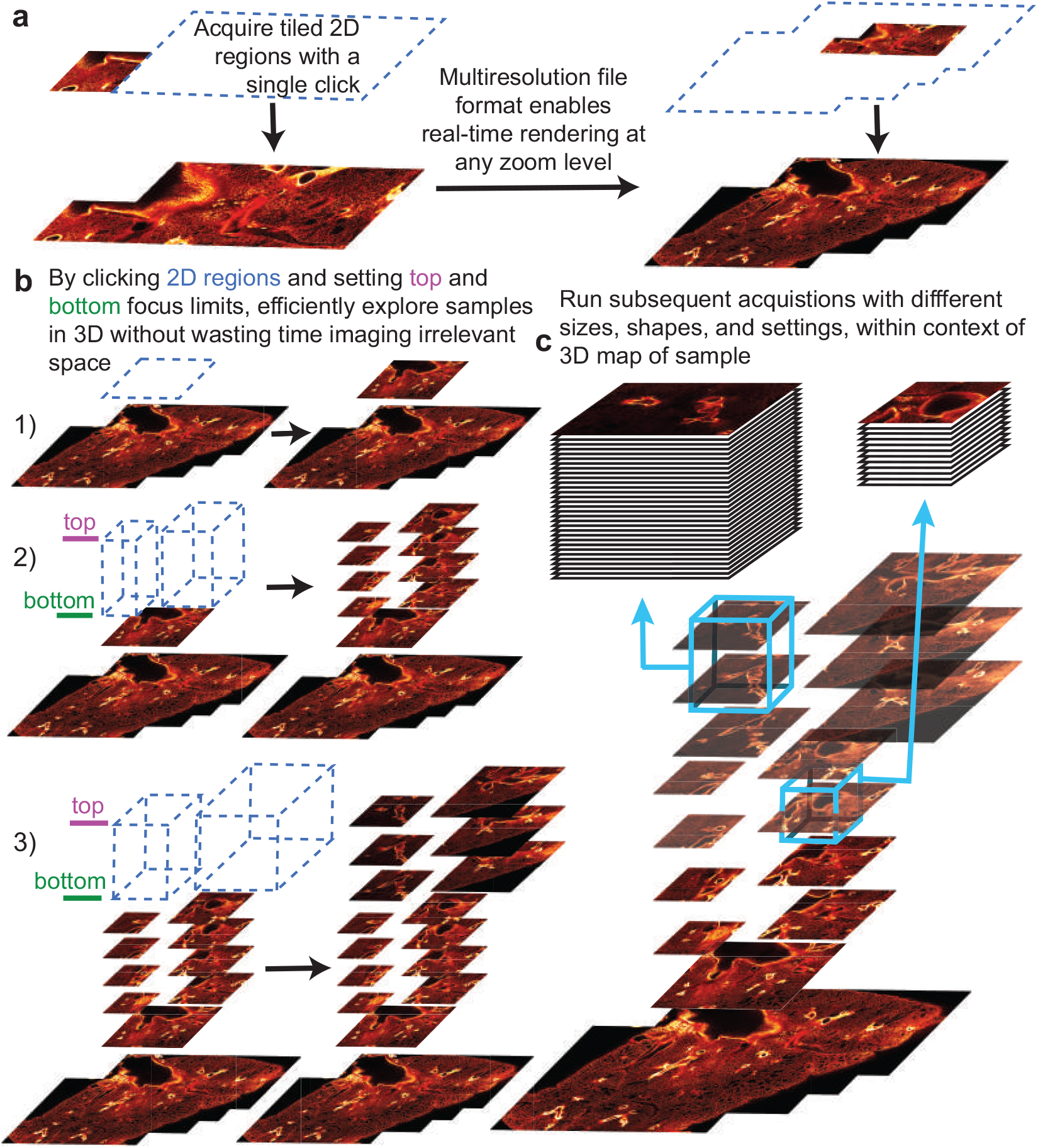
Explore acquisitions. (A) A simple and intuitive user interface enables panning, zooming, and acquisition of high-resolution images by tiling together multiple fields of view (B) Map complex structures in 3D (a branching lung airway shown here) by setting upper and lower focus limits to collect z-stacks over arbitrarily chosen 2D regions (C) Use the coarse-grained sample mapping to run subsequent acquisitions within the context of the greater sample morphology (high resolution z-stacks at airway branch points shown here)

Once samples have been mapped using explore acquisitions, μMagellan allows the creation of arbitrarily shaped 2D and 3D ROIs in the coordinate space of specimens using interpolated surfaces. These ROIs speed up acquisitions because they allow users to quickly specify non-cuboidal volumes for imaging, avoiding time-consuming acquisition of irrelevant space. Importantly, they can also be used to automatically vary hardware settings during acquisition based on sample morphology using covariant pairings (described below). Users mark points on 2D transverse slices **(Fig. 2a,b),** which are then used to interpolate 3D surfaces using an algorithm based on Delaunay triangulations^13^ **(Fig. 2c,d, Supplementary note 3).** Acquisitions can thus be run over 3D ROIs based on specified distances above and below a planar or non-planar surface. For example, the former could be used to collect the full volume of a slide-mounted sample that isn’t orthogonal to the optical axis. The latter could be used to specify the exact volume extending from the surface of a rounded organ to the depth limit of a particular microscope. Alternatively, more complex volumes can be created combining two surfaces to bound arbitrarily shaped ROIs **(Fig. 2e).**

**Figure 2.**
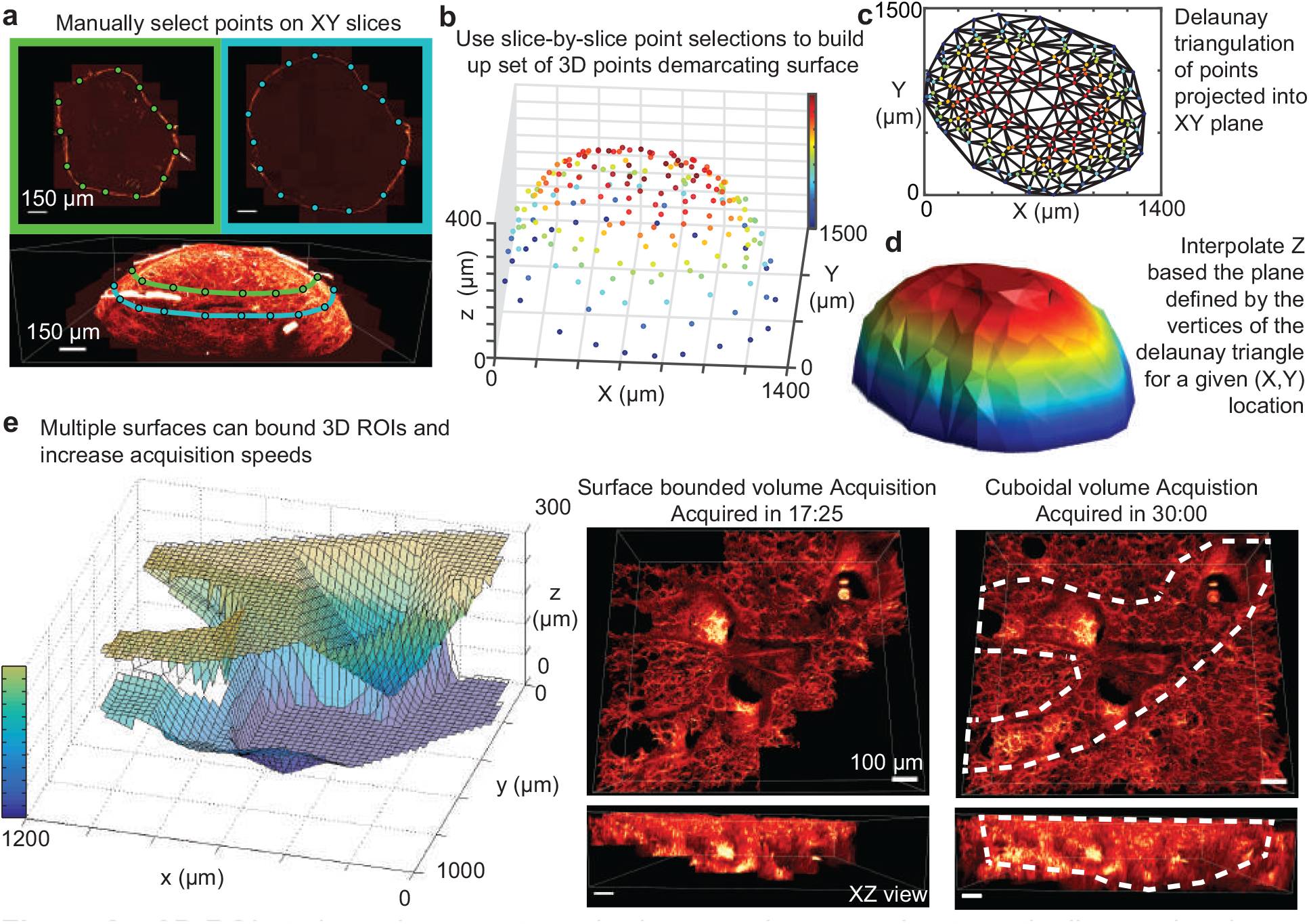
3D ROIs to bound non-rectangular image volumes and automatically vary hardware settings during acquisition. **(A)** Interpolation points are marked on 2D slices of a popliteal lymph node **(B)** A full set of 3D points for interpolation **(C)** Delaunay triangulation of points projected onto XY plane **(D)** A fully interpolated 3D surface used to automatically run a non-rectangular acquisition **(E)** Two surfaces used to bound an airway in a 400 urn lung slice, which significantly reduces acquisition time compared to acquiring the entire bounding cuboidal volume

To provide automation necessary for complex experiments and facilitate reproducible experiments across specimens, μMagellan exposes all hardware settings to automated variation through the concept of covariant pairings. A covariant pairing allows the specification of a particular hardware setting (such as excitation power, detector gain, exposure, etc.) based on either another hardware setting or, as related to sample morphology, based on a calculation involving the geometry of a particular surface. Users specify pairs of values between covariants, and prior to the acquisition of each image μMagellan sets the value of the dependent based on the value of the independent, linearly interpolating if both are numerical. By associating variation in settings with sample morphology, settings can be easily adapted to samples and reused within and between experiments to facilitate comparisons **(Supplementary Fig 1).**

μMagellan also has several features that optimize the imaging of living specimens across long time scales. The μMagellan acquisition engine is designed for flexibility, and almost all settings (such as spatial regions, time point spacing, automated excitation calculations, etc.) can be altered during acquisition. This flexibility enables the ability to adapt to dynamic biological processes and investigate behaviors at a variety of scales of space and time within a single experiment.

Additionally, μMagellan provides integrated on-the-fly changes that accommodate drift or morphological changes, which cause samples to move out of view of the imaging volume and the delivery of inappropriate amounts of excitation power **(Supplementary Fig. 2a).** This is particularly important in the case of multiphoton and confocal microscopy, which have been used to image samples for long periods of time (24 hours or more) without compromising viability^14,15^ but in which sample drift often besets automated acquisition. In addition to allowing for manual adjustments to these parameters during experiments, μMagellan provides an algorithm for automated drift compensation based on the 3D cross correlation between images from a designated fiduciary channel over the previous 2 time points.^16^ The maximum value of the cross correlation gives an estimate of the vertical drift **(Supplementary Fig. 2b),** which can be used to adjust a focus drive to offset this drift **(Supplementary Fig. 2c)** and provide a correctly registered image volume indefinitely **(Supplementary Fig. 2d).**

Finally, μMagellan provides automation to run multiple acquisitions in series or in parallel. For example, a slide containing multiple samples can be explored, and the locations of each sample identified with the creation of surfaces. Subsequently, acquisitions can be set for each sample and sequentially imaged in an automatic fashion for hours or days at a time. Parallel acquisitions can be used to monitor multiple sites in a sample simultaneously or multiple organs from the same animal to compare conditions while minimizing biological variability.

As new methods continue to be developed in order to probe the structure, function, and dynamics of biological systems in new ways, powerful and flexible software to put these tools into practice are becoming increasingly important. μMagellan fills an important niche in the open-source bioimaging software ecosystem by empowering many existing microscopes for automated, reproducible, high throughput applications. Its open source code also makes it an ideal platform for the development and dissemination of new technologies, thereby increasing the ease with which they can be put into practice to reveal the mysteries of biological systems. μMagellan comes bundled with μMagellan and can be accessed in the plugins menu under the “Acquisition tools” group. Documentation for installing and using μMagellan can be found within the software itself and at https://micro-manager.org/wiki/MicroMagellan.

## ACKNOWLEDGMENTS

We thank K. Thorn, M. Tsuchida, and C. Weisiger for helpful conversations during development, M. Headley for his feedback and beta testing, and E. Oswald for providing the lung data used in figure 1. This work was supported in part by US National Institutes of Health grants R01 AI52116 and U19A1077439-06 and the Sandler Basic Asthma Research Center.

## AUTHOR CONTRIBUTIONS

H.P. and M.F.K. conceived of the project. H.P. designed and programmed the software with help from N.S. H.P. and K.C. performed beta testing. H.P., K.C. and M.F.K. wrote the manuscript. K.C. wrote the user guide. M.F.K. and R.V. provided administrative and financial support.

## COMPETING FINANCIAL INTERESTS

N.S. and R.V. are co-founders of Open Imaging, the company that maintains and develops μMagellan.

**Supplementary Figure 1.**
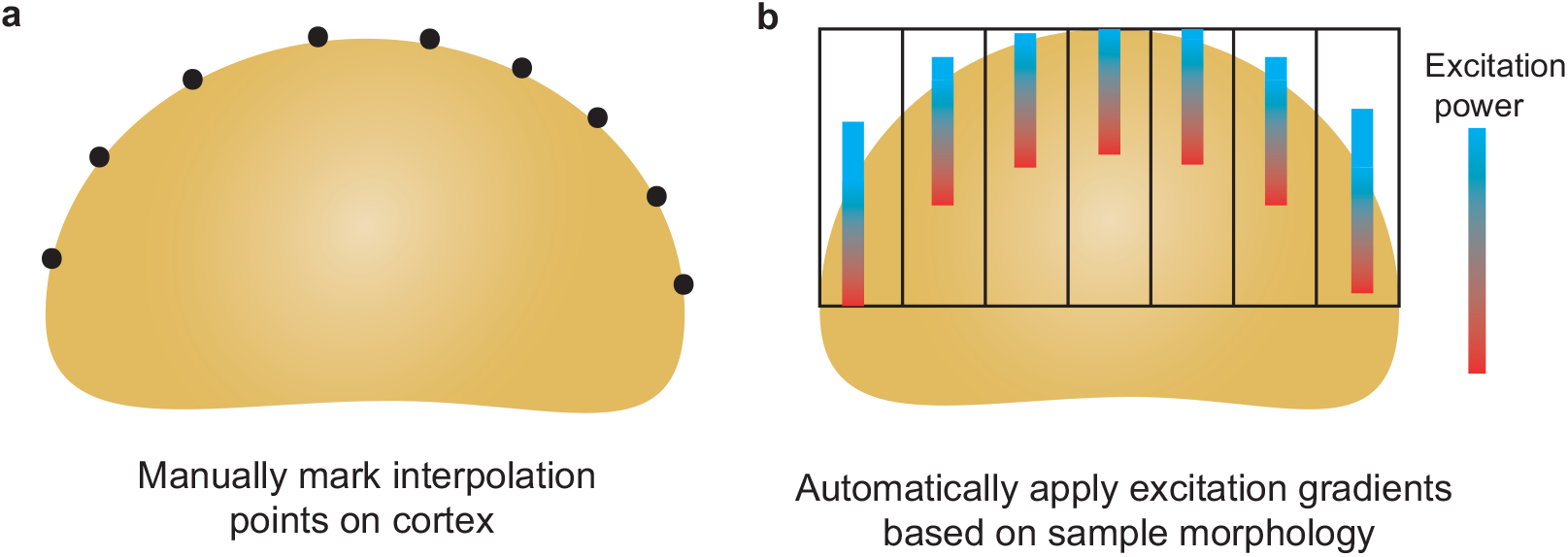
Surfaces can be used to automatically vary acquisition parameters based on sample morphology. **(A)** Interpolation points on lymph node cortex define surface **(B)** Gradients of increasing excitation power at different XY positions, which begin at the top of the sample for each position.

**Supplementary Figure 2.**
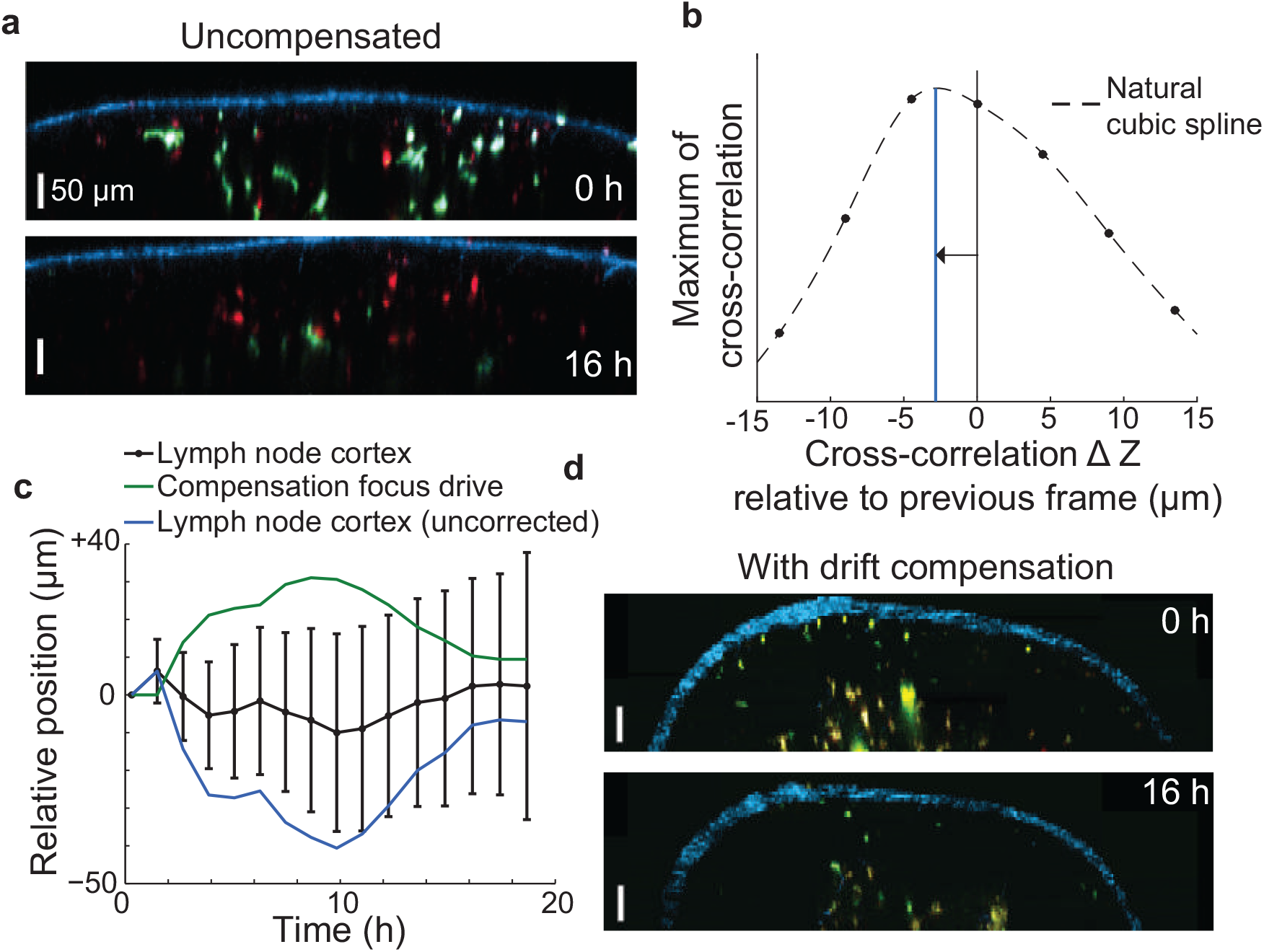
Automatic 3D cross correlation based drift compensation. (A) XZ slice of popliteal lymph node explant imaging showing sample drift between start and hour 16 of imaging. (B) Maximum value of 3D cross correlation between second harmonic generation channel of two successive time points, with a 3D cubic spline fit to allow for sub-voxel accuracy. (C) Applying a focus offset based on (B) compensates for sample drift. Position of lymph node cortex calculated as mean of 15 reference points. Error bars show their standard deviation. (D) Drift compensation allows stabilized imaging of lymph node over long time course

**Supplementary Note 1. Calibration and stitching**

In order to accurately stitch together images from multiple XY positions, μMagellan must be calibrated to determine the affine transformation matrix that relates image pixel coordinates to those of the XY stage. This calibration must be performed separately for each pixel size setting defined within μMagellan. The “Calibrate” button on the bottom right corner of the μMagellan GUI provides manual and automatic means of doing so. The automatic method prompts user’s to take three images, moving the XY stage to a different location for each one, then calculates the cross-correlation of these images relative to one another in order to calculate the entries of the affine transform. The manual method allows user’s to input specific values for rotation, shear, x scale, and y scale, and is particularly useful for making small adjustments if an automatic calibration is slightly off. This can be easily accomplished by making adjustments, pressing apply, running a 2x2 field of view acquisition to determine how fields of view are aligned, and repeating.

**Supplementary Note 2. File Format**

μMagellan saves data in a mipmapped (multi-resolution), Tiff-based format. Inside the root saving directory, there is directory titled “Full resolution”, an arbitrary number of directories titled “Downsampled_x#” where the #’s are successive powers of 2, and an XML file that allows reading by the BigDataViewer.^3^ Within each of these directories are a series of Tiff stacks, with each Tiff stack corresponding to all channels/z slices/time points at a given XY position (tile). If the data at a given tile exceeds 4 GB, images are distributed across multiple Tiff stacks with a number appended to the end of the filename. When viewed by Magellan or the BigDataViewer, tiles are dynamically stitched together to form a contiguous image. However, storing them individually facilitates custom post-acquisition processing pipelines such as flat-field correction/background subtraction or registration based stitching.

At each successively lower resolution, a 2x2 grid of images from the previous level is binned by a factor of two to create a lower resolution image with the same pixel dimensions. This downsampling scheme allows random access to image data by ensuring that the same number of pixels is read from disk regardless of the requested zoom level.

Within each Tiff stack, a map of the locations of individual images is written to facilitate fast reading without having to scan entire files. Custom processing pipelines can make use of this feature to quickly read data by instantiating the MultiResMultipageTiffStorge class, which can be found within the Magellan.jar file that is bundled with μMagellan.

**Supplementary Note 3. Interpolation algorithm**

μMagellan allows the generation of arbitrarily shaped 3D surfaces based on interpolation points marked on 2D slices. Users mark interpolation points on XY slices corresponding to different locations along the Z axis **(Fig. 1a),** in order to build up a set of XYZ interpolation points **(Fig. 1b).** These XYZ points are then projected onto the XY plane, and their Delaunay triangulation is calculated **(Fig. 2c).** Z values can then be calculated as a function of (X,Y) points by finding the Z coordinate on the plane specified by the three (XYZ) vertices of the Delaunay triangle enclosing a given point on the 2D (XY) projection. In this way a full 3D surface can be interpolated **(Fig. 2d).** Importantly, there is only a single Z value for each XY location using this interpolation scheme. More complex volumes can be created by using multiple surfaces to bound 3D volumes for acquisition **(Fig. 2e).** In order to speed up calculations that utilize surface geometry during acquisition, look up tables (LUTs) of surface interpolation Z values as a function of (X,Y) are automatically calculated each time an interpolation point is added or removed. Projections of the parts of a surface interpolation that intersect a given 2D slice can be shown overlaid on image data, allowing users to visualize surfaces relative to samples. To increase the speed of this process and allow for real time user feedback, these LUTs are calculated and displayed first at low resolution, followed by increasingly high levels of detail.

